# Bimodal distribution of coral bleaching prevalence is consistent with state transition dynamics in thermal stress response: five years of standardised monitoring in Japan

**DOI:** 10.64898/2026.03.05.709742

**Authors:** Hiroki Fukui

## Abstract

Coral bleaching is conventionally modelled as a continuous response to cumulative thermal stress, yet the distributional structure of site-level bleaching prevalence has rarely been examined. Here we analyse five years (fiscal years 2020–2024) of standardised bleaching surveys from Japan’s Monitoring Site 1000 program, encompassing 2,288 observations across 585 survey points at 26 sites in the Ryukyu Archipelago and adjacent waters. We document three principal findings. First, bleaching prevalence is bimodally distributed in all five years: the intermediate range (20–80%) remains stable at 21–27% of observations, while inter-annual variation is driven by redistribution between low and high domains — confirmed by Hartigan’s dip test (all p < 0.001) and beta mixture modelling (ΔBIC = 9.0–113.9). Second, the 2022 and 2024 bleaching events are qualitatively distinct: 2022 was a partial mass bleaching (positively skewed, selective), while 2024 was comprehensive (symmetric, median 60.0%). Third, a simple threshold metric (days above 30°C) outperformed Degree Heating Weeks in discriminating bleaching across all severity levels (GEE-based AUC: 0.877 vs 0.624 at ≥50% prevalence, p < 0.001; 0.830 vs 0.633 at >0%, p = 0.007), indicating that metric structure and the ecological severity threshold defining the outcome are inseparable design considerations.

## 1.1 1. Introduction

Mass coral bleaching, driven by marine heatwaves of increasing frequency and severity, has become the dominant threat to tropical and subtropical reef ecosystems worldwide (Hughes et al. 2018a). The standard predictive framework centres on Degree Heating Weeks (DHW), a cumulative thermal stress metric that integrates the magnitude and duration of sea surface temperatures exceeding a site-specific climatological maximum (Liu et al. 2014). The logic of DHW is inherently cumulative: thermal stress is treated as a dose that accumulates continuously, and bleaching is predicted to intensify as a graded function of that dose. DHW thresholds of 4 and 8 °C-weeks are operationally interpreted as indicating the onset and severe phases of mass bleaching, respectively. But this cumulative logic carries two assumptions that have received limited empirical scrutiny at the site level. The first is linearity: bleaching prevalence should increase progressively with heat exposure. The second is universality: standard thresholds are applied globally, although regional studies have demonstrated substantial variation in effective thresholds (Sakai et al. 2019; Whitaker & DeCarlo 2024) and in the relative performance of alternative metric formulations (Lachs et al. 2021). If instead bleaching is a state transition — a threshold-mediated switch between an unbleached and a bleached state — then cumulative heat dose may not be the most appropriate predictor category for ecologically significant bleaching.

The distributional structure of bleaching prevalence — how site-level values are distributed across a monitoring network — provides a direct test of the linearity assumption. If bleaching is a continuous dose-response, one would expect prevalence values to be unimodally distributed, spreading progressively with increasing thermal stress. If instead bleaching involves a threshold-mediated state transition, as suggested by bioenergetic models of the photodamage–symbiont expulsion cascade (Pfab et al. 2024), one would predict a bimodal distribution: sites cluster into low and high bleaching states with few observations at intermediate values, and inter-annual variation is expressed as redistribution between states rather than as a shift along a continuum.

Examining whether the distributional data are consistent with this prediction requires a dataset with three properties: continuous (not binary) measurement of bleaching prevalence at the site level, standardised protocols across sites, and repeated observations over multiple years encompassing both mild and severe bleaching events. Japan’s Monitoring Site 1000 coral reef survey program meets all three criteria. Administered by the Ministry of Environment, the program monitors 26 sites spanning approximately 9° of latitude, from the Yaeyama Islands (24.3°N) to Kushimoto (33.5°N) — the northern distribution limit of hermatypic corals in the northwestern Pacific. Bleaching prevalence is recorded as a continuous percentage at each of 585 survey points, and the five fiscal years 2020–2024 encompass two major bleaching events (2022 and 2024) bracketing three milder years. Despite more than two decades of continuous monitoring since 2003, the Monitoring Site 1000 coral reef dataset has not been the subject of an international peer-reviewed study (Kawagoe 2017 reported 2016 bleaching in the journal of the Japanese Coral Reef Society; Kimura et al. 2022 incorporated the data into an ICRI regional report). The present analysis represents the first such use.

In this study, we examine the distributional structure of bleaching prevalence across this network and report three findings. First, bleaching prevalence is bimodally distributed in all five years, with inter-annual variation driven by redistribution between low and high domains. Second, the 2022 and 2024 bleaching events are qualitatively distinct — partial versus comprehensive mass bleaching. Third, a simple threshold exceedance metric (days above 30°C) discriminates bleaching more effectively than DHW across all severity levels, with the largest advantage at moderate-to-severe thresholds — indicating that metric structure and response definition are inseparable design considerations.

## 1.2 2. Methods

### 1.2.1 2.1 Study area and data source

We analysed coral bleaching data from the Monitoring Site 1000 coral reef survey program, administered by the Ministry of Environment of Japan (Ministry of Environment 2024). The program monitors coral reef condition across the Ryukyu Archipelago and adjacent areas using a standardised spot-check protocol. The dataset used in this study covers fiscal years 2020–2024 (data citation: Ministry of Environment, Japan. Monitoring Site 1000 Coral Reef Survey (FY2020–2024); approval number Kansei-Tahatsu No. 2602271) and comprises 585 survey points across 26 sites, yielding 2,288 observations.

Survey sites span approximately 9° of latitude, from the Yaeyama Islands (24.3°N) to Kushimoto (33.5°N), encompassing the northern distribution limit of hermatypic corals in the northwestern Pacific. Sites are structured as clusters of survey points (typically 10–40 points per site) within a ∼5 km^2^ area.

### 1.2.2 2.2 Bleaching and mortality assessment

At each survey point, bleaching prevalence (bleaching_all) was recorded as the percentage of colony area showing signs of bleaching, averaged across quadrats within the point. Mortality prevalence (mortality_all) was recorded using the same protocol. Both variables yield values at approximately 0.5% resolution (135 unique values across the dataset), reflecting the original recording protocol. Surveys are conducted annually in autumn (September–November), approximately 1–3 months after the summer thermal stress peak. Consequently, the mortality values recorded represent the state at the time of survey and may underestimate total post-bleaching mortality, which can progress over six months or longer (Sakai et al. 2019). This temporal constraint applies to all mortality results reported in this study.

### 1.2.3 2.3 Sample size variation

The number of survey sites ranged from 21 to 25 per year, with 428–496 survey points per year (Table 1). Variation was driven primarily by the addition or removal of entire sites: three sites (02, 08, 20) were added in FY2022, none of these three were surveyed in FY2023 (Table 1), and site 25 was added in FY2024. Minor variation arose from changes in point numbers within sites (e.g., site 05 ranged from 32 to 42 points). Missing data were negligible (1 record in FY2024). Twenty-one of 26 sites participated in all five fiscal years; sensitivity analyses restricted to this balanced panel are reported in the Supplementary Material (Fig. S1).

**Table 1.** Sample sizes by fiscal year.

| Fiscal year | Sites | Survey points |
| --- | --- | --- |
| 2020 | 21 | 428 |
| 2021 | 22 | 438 |
| 2022 | 25 | 496 |
| 2023 | 22 | 455 |
| 2024 | 23 | 471 |
| Total | 26 | 2,288 |

### 1.2.4 2.4 Satellite sea surface temperature

Daily sea surface temperature (SST) was obtained from the NASA MUR SST v4.1 product (Multi-scale Ultra-high Resolution SST; JPL MUR MEaSUREs Project 2015), which provides global coverage at 0.01° (∼1 km) resolution. SST values were extracted for each monitoring site using the nearest grid point to the site centroid.

## 1.2.5 2.5 Thermal stress metrics

Two thermal stress metrics were computed from the same SST product (MUR SST v4.1), so that the comparison between them reflects the difference in metric structure rather than differences in input data or spatial resolution.

**Degree Heating Weeks (DHW)** were calculated following the standard NOAA Coral Reef Watch methodology (Liu et al. 2014). The Maximum Monthly Mean (MMM) climatology was calculated as the highest monthly mean SST over the 2003–2014 baseline period — the longest span of complete annual cycles available in MUR at the time of analysis (MUR v4.1 production begins mid-2002). HotSpots were defined as max(SST ™ MMM, 0), accumulated over an 84-day (12-week) window, and divided by 7 to yield DHW in °C-weeks. This implementation differs from the operational NOAA CRW algorithm in one respect: the standard CRW algorithm applies a HotSpot > 1°C cutoff, accumulating only anomalies exceeding MMM + 1°C (Liu et al. 2014), whereas our implementation accumulates all positive anomalies above MMM. This choice follows the finding by Lachs et al. (2021) that HotSpot thresholds at or below MMM improve predictive accuracy. **Threshold exceedance days** were defined as the number of days with MUR SST exceeding 30°C over the summer period preceding each survey. This threshold was adopted a priori based on Sakai et al. (2019), who identified 30°C as associated with bleaching onset at Sesoko Island, Okinawa.

## 1.2.6 2.6 Statistical analyses

### 1.2.6.1 Distributional analysis

Violin plots of bleaching prevalence and mortality prevalence were generated using Gaussian kernel density estimation with Scott’s rule bandwidth selection (scipy.stats.gaussian_kde; Scott 1992). Box plots showing interquartile range and median are overlaid within each violin.

### 1.2.6.2 Tests for bimodality

Bimodality of bleaching prevalence was assessed using two independent approaches. First, Hartigan’s dip test (Hartigan & Hartigan 1985) was applied to the full data (0–100%) and to the intermediate range (0 < bleaching < 100%) separately, to confirm that bimodality is not an artefact of boundary inflation at 0% and 100%.

Second, one-component and two-component beta mixture models were fitted to the intermediate-range data (0 < bleaching < 100%, scaled to [0, 1]) using an expectation-maximisation (EM) algorithm with 50 random initial parameter sets (3-set and 50-set runs yielded identical BIC values; convergence criterion: ΔBIC < 1e-6 between iterations, maximum 1000 iterations). Model selection was based on the Bayesian Information Criterion (BIC), with ΔBIC > 10 indicating very strong support for the more complex model (Kass & Raftery 1995).

### 1.2.6.3 Intraclass correlation

Intraclass correlation coefficients (ICC) were estimated from a one-way random-effects ANOVA model with site as the grouping factor, using bleaching prevalence in percentage scale. Results were robust to logit-transformed (0, 1) scaling with ε-adjustment for boundary values.

### 1.2.6.4 Predictive comparison

The predictive accuracy of DHW and threshold exceedance days for bleaching was compared using ROC analysis and the area under the curve (AUC). Both metrics were derived from the same SST product (MUR), isolating the effect of metric structure on predictive performance. Because satellite SST was available for 21 of 26 monitoring sites (five sites lacked MUR coverage), the predictive comparison was conducted on these 21 sites across all five fiscal years (n = 2,142 observations). Three binary response definitions were used: (1) bleaching occurrence (bleaching_all > 0%), (2) moderate bleaching (bleaching_all ≥ 50%), and (3) severe bleaching (bleaching_all ≥ 80%). This multi-threshold approach tests whether predictive performance depends on the ecological severity of the response. To address spatial non-independence of observations within sites, we applied generalised estimating equations (GEE) with an exchangeable correlation structure (statsmodels v0.14, Python), clustering observations by site (n = 21 sites). This approach accounts for within-site correlation without requiring strong parametric assumptions about the random-effects structure. GEE-based AUC values were computed as the AUC of GEE-predicted probabilities. Pairwise comparison of AUC values between metrics was performed using a cluster bootstrap Wald test (10,000 iterations, site-level resampling, seed = 42), which preserves the within-site correlation structure that conventional tests such as DeLong (1988) do not accommodate.

The analyses are descriptive and exploratory. The 30°C threshold was adopted a priori from Sakai et al. (2019); all other thresholds (27–32°C) were examined as robustness checks. No correction for multiple comparisons was applied; results should be interpreted as hypothesis-generating.

### 1.2.6.5 Conditional mortality analysis

Mean and median mortality were computed for five bleaching prevalence bands (0%, 0–20%, 20–50%, 50–80%, ≥80%) to characterise the bleaching–mortality relationship.

### 1.2.6.6 Spot-level tracking

Survey points with data in consecutive years were tracked to quantify recovery and cumulative impact trajectories. Specifically, 439 points with data in both 2022 and 2023 were used for recovery analysis, and 435 points with data in both 2022 and 2024 were used for back-to-back bleaching analysis.

All analyses were conducted in Python 3.11 using pandas, scipy, numpy, and matplotlib. Code is available at https://github.com/hirokifukui/marine-obs

## 1.3 3. Results

### 1.3.1 3.1 Bleaching prevalence across five years

Bleaching prevalence varied substantially across the five-year study period (Fig. 1, Table 2). The years 2020, 2021, and 2023 showed low overall bleaching, with median prevalence of 0% and 38–49% of survey points recording any bleaching. In contrast, 2022 and 2024 were major bleaching years, with 78.0% and 85.1% of points recording bleaching, respectively. The year 2024 was the most severe, with a median bleaching prevalence of 60.0% and a mean of 48.4%.

**Fig. 1.** Annual distribution of coral bleaching prevalence (upper panel) and mortality prevalence (lower panel) across the Monitoring Site 1000 network, fiscal years 2020–2024. Violin plots show kernel density estimates (Gaussian kernel, Scott’s rule bandwidth); KDE tails may extend slightly beyond the 0–100% data bounds. Box plots within each violin show median (horizontal line) and interquartile range. The percentage of survey points with bleaching > 0% (upper panel) or mortality > 0% (lower panel) is annotated above each year. Sample sizes: n = 428, 438, 496, 455, 471 for FY2020–2024, respectively.

**Table 2.** Bleaching prevalence (bleaching_all, %) by fiscal year.

| Year | n | Median | Mean | Q25 | Q75 | >0% |
| --- | --- | --- | --- | --- | --- | --- |
| 2020 | 428 | 0.0 | 17.0 | 0.0 | 27.5 | 49.3% |
| 2021 | 438 | 0.0 | 16.8 | 0.0 | 32.1 | 38.4% |
| 2022 | 496 | 20.0 | 37.6 | 0.5 | 80.0 | 78.0% |
| 2023 | 455 | 0.0 | 12.3 | 0.0 | 23.8 | 44.8% |
| 2024 | 471 | 60.0 | 48.4 | 2.5 | 90.0 | 85.1% |

### 1.3.2 3.2 Bimodal distribution of bleaching prevalence

The violin plots revealed a bimodal distribution in all five years (Fig. 1). Survey points clustered into a low-bleaching domain (<20%) and a high-bleaching domain (≥80%), with the intermediate range (20–80%) remaining stable at 21–27% of observations regardless of year (Table 3). Inter-annual variation was driven by redistribution between the low and high domains: in 2024, the high domain contained 39.4% of observations, compared to 2.7% in 2021.

**Table 3.**
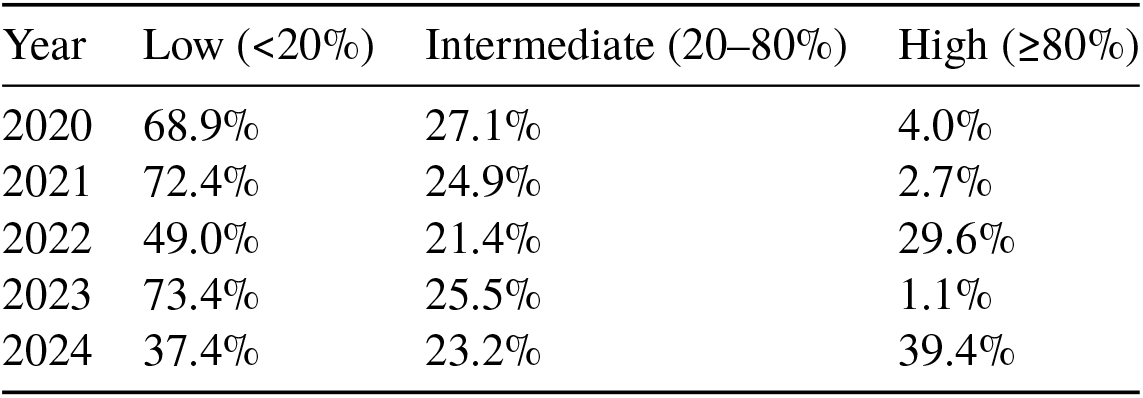
Distribution of survey points across bleaching domains.

| Year | Low (<20%) | Intermediate (20–80%) | High (≥80%) |
| --- | --- | --- | --- |
| 2020 | 68.9% | 27.1% | 4.0% |
| 2021 | 72.4% | 24.9% | 2.7% |
| 2022 | 49.0% | 21.4% | 29.6% |
| 2023 | 73.4% | 25.5% | 1.1% |
| 2024 | 37.4% | 23.2% | 39.4% |

Hartigan’s dip test rejected the null hypothesis of unimodality in all years, both for the full data and for the intermediate range (0 < bleaching < 100%; all p < 0.001; Table 4). The dip statistic was largest in 2024 (D = 0.132, full data), consistent with the most pronounced bimodality in the most severe bleaching year.

**Table 4.** Hartigan’s dip test results.

| Year | Full data D | Full data p | Intermediate D | Intermediate p |
| --- | --- | --- | --- | --- |
| 2020 | 0.044 | <0.001 | 0.055 | <0.001 |
| 2021 | 0.037 | <0.001 | 0.097 | <0.001 |
| 2022 | 0.091 | <0.001 | 0.116 | <0.001 |
| 2023 | 0.055 | <0.001 | 0.063 | <0.001 |
| 2024 | 0.132 | <0.001 | 0.105 | <0.001 |

Two-component beta mixture models were favoured over one-component models in every year (ΔBIC = 9.0–113.9; Table 5). Support was very strong (ΔBIC > 10) in four of five years; in 2023, ΔBIC = 9.0 represents strong evidence (Kass & Raftery 1995), but the qualitative pattern — a low-bleaching component and a high-bleaching component — was consistent across all years including 2023. In 2023 specifically, the weaker statistical support reflects a year of low overall bleaching in which the high-bleaching component was small (w = 0.299) and its parameters overlapped more with the low-bleaching component than in other years; the qualitative two-state structure is nonetheless present. In 2023, the high-bleaching component parameters (α = 8.37, β = 10.81) describe a nearly symmetric distribution centred below 0.5 — qualitatively distinct from the right-skewed high-bleaching components in other years (e.g., 2024: α = 6.17, β = 1.49). This suggests that in mild years, the two-state structure represents a low-bleaching state and a moderate-bleaching state rather than the low/high polarisation observed in major bleaching years. The two components consistently showed a low-bleaching group (α < 1, β > 2; J-shaped) and a high-bleaching group (α > β in major bleaching years; right-skewed).

**Table 5.** Beta mixture model comparison (intermediate range, 0 < bleaching < 100%). ΔBIC > 10 indicates very strong support for the two-component model (Kass & Raftery 1995); 2023 (ΔBIC = 9.0) represents strong evidence.

| Year | n | BIC | BIC | $\Delta\text{BIC}$ | w | $\alpha$ | $\beta$ | w | $\alpha$ | $\beta$ |
| --- | --- | --- | --- | --- | --- | --- | --- | --- | --- | --- |
| 2020 | 211 | −82.4 | −113.7 | 31.4 | 0.740 | 0.69 | 3.07 | 0.260 | 23.66 | 7.41 |
| 2021 | 168 | −24.0 | −79.1 | 55.2 | 0.396 | 0.47 | 2.94 | 0.604 | 6.69 | 3.56 |
| 2022 | 379 | −120.7 | −224.3 | 103.6 | 0.503 | 0.75 | 5.47 | 0.497 | 7.14 | 1.42 |
| 2023 | 204 | −148.1 | −157.0 | 9.0 | 0.701 | 0.56 | 2.06 | 0.299 | 8.37 | 10.81 |
| 2024 | 381 | −85.3 | −199.2 | 113.9 | 0.384 | 0.60 | 4.74 | 0.616 | 6.17 | 1.49 |

### 1.3.3 3.3 Qualitative distinction between the 2022 and 2024 bleaching events

Although both 2022 and 2024 were major bleaching years, their distributional signatures differed (Fig. 1). In 2022, the distribution was positively skewed (skewness = 0.413), with most points in the low-to-moderate range and a tail of severe cases. In 2024, the distribution was approximately symmetric (skewness = −0.100), with the median at 60.0% and the high-bleaching component of the beta mixture model dominant (w = 0.616 vs 0.497 in 2022).

### 1.3.4 3.4 Non-linear relationship between bleaching and mortality

Mortality prevalence was structurally distinct from bleaching prevalence: right-skewed rather than bimodal, with the bulk of observations below 20% even in severe bleaching years (Table 6).

**Table 6.**
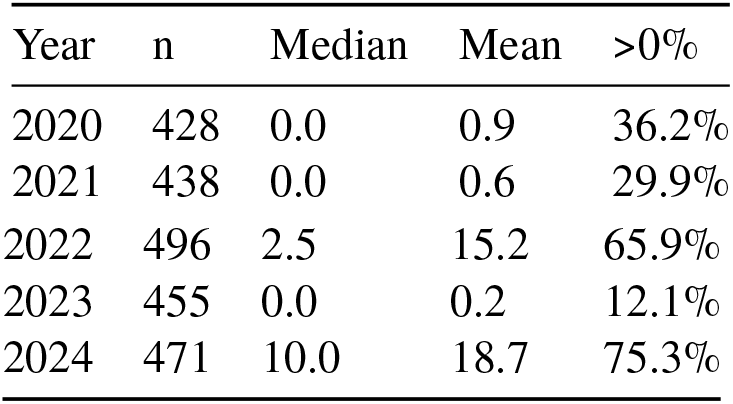
Mortality prevalence (mortality_all, %) by fiscal year.

Conditional mortality analysis revealed a non-linear relationship with bleaching prevalence (Table 7). Mortality escalated sharply above 80% bleaching, yet even at 80–100% bleaching, the median mortality was only 25.0%.

**Table 7.** Conditional mortality by bleaching prevalence band (pooled 2020–2024).

| Bleaching band | n | Mortality mean | Mortality median | Mortality >0% |
| --- | --- | --- | --- | --- |
| 0% | 917 | 0.0 | 0.0 | 1.1% |
| 0–20% | 448 | 1.5 | 0.5 | 50.9% |
| 20–50% | 271 | 4.8 | 1.0 | 69.7% |
| 50–80% | 285 | 13.2 | 2.0 | 84.9% |
| ≥80% | 366 | 30.9 | 25.0 | 96.4% |

### 1.3.5 3.5 Spatial structure of bimodality

The bimodal distribution was driven primarily by between-site divergence rather than within-site variability. In 2024, six sites (11, 12, 18, 19, 24, 25) had all points in the low domain (<20% bleaching), while seven sites (6, 10, 13, 14, 15, 16, 17) had all points in the high domain (≥80%). Intraclass correlation coefficients (ICC) confirmed this spatial partitioning: site identity explained 76–88% of the variance in bleaching prevalence across years (Table 8). ICC was highest in 2021 (0.887), a mild year in which site-level differences in baseline vulnerability were most apparent, and lowest in 2024 (0.760), when the comprehensive event partially overrode site-level variation.

**Table 8.**
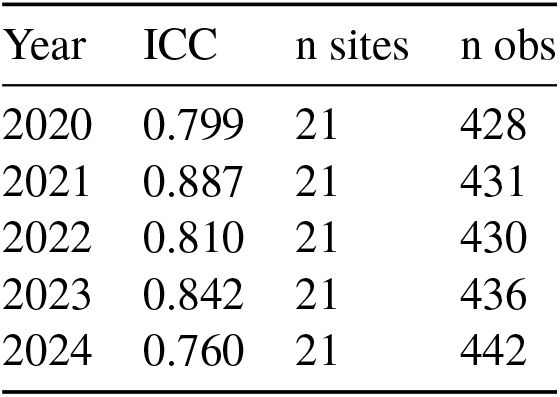
Intraclass correlation coefficients for bleaching prevalence by fiscal year.

ICC computed on the balanced panel of 21 sites with data in all five fiscal years (sites 02, 08, 20, 25, 26 excluded). n obs: survey points at these 21 sites with valid bleaching records in each year.

### 1.3.6 3.6 Recovery and back-to-back bleaching

Bleaching prevalence declined sharply from 2022 (mean 37.6%) to 2023 (mean 12.3%). Among 192 survey points with ≥50% bleaching in 2022, 28.1% recorded zero bleaching in 2023, and 85.4% showed zero mortality at the 2023 survey.

Tracking 435 points across 2022 and 2024 revealed three trajectories: 32.0% experienced ≥50% bleaching in both years; 9.4% transitioned from 0% in 2022 to ≥50% in 2024; 5.5% showed the reverse (≥50% to 0%).

### 1.3.7 3.7 Predictive comparison: threshold exceedance versus DHW

The predictive comparison was conducted on the 21 sites with satellite SST coverage across all five fiscal years (n = 2,142 observations; see Section 2.6). Days above 30°C significantly outperformed DHW across all three response definitions (Table 9). For bleaching occurrence at the detection limit (>0%), days above 30°C was superior (AUC 0.830 vs 0.633; GEE cluster bootstrap Wald test, z = 2.71, p = 0.007). For moderate bleaching (≥50%), the advantage was most pronounced (AUC 0.877 vs 0.624; z = 4.51, p < 0.001). For severe bleaching (≥80%), days above 30°C again outperformed DHW (AUC 0.895 vs 0.696; z = 3.08, p = 0.002).

**Table 9.** GEE-based AUC values for days above 30°C and DHW across three bleaching response thresholds. All values derived from generalised estimating equations (GEE) with exchangeable correlation structure and site-level clustering (GEE fitted to 2,142 survey-point observations clustered by 21 sites (93 site-years with complete predictor data)). Within-site correlation (ρ) is reported for each metric; higher ρ for DHW indicates greater dependence on site-level fixed effects. Bootstrap 95% CI (10,000 iterations, site-level resampling) are reported in Supplementary Table S4

| Response threshold | Days above $30^{\circ}\text{C}$ AUC | DHW AUC | Within-site $\rho$ (days30 / DHW) |
| --- | --- | --- | --- |
| $>0\%$ (any bleaching) | 0.830 | 0.633 | 0.172 / 0.339 |
| $\geq 50\%$ (moderate–severe) | 0.877 | 0.624 | 0.110 / 0.261 |
| $\geq 80\%$ (severe) | 0.895 | 0.696 | 0.036 / 0.205 |

Both metrics were derived from the same SST product, so differences in AUC reflect metric structure rather than input data quality. Notably, DHW’s AUC was similar across the >0% and ≥50% response definitions (0.633 and 0.624), while the threshold metric’s AUC increased sharply from 0.830 to 0.877 — consistent with threshold exceedance logic capturing a binary state transition that becomes ecologically manifest only above a minimum severity. The lower within-cluster correlation for the days above 30°C model (ρ = 0.110 at ≥50%) compared to DHW (ρ = 0.261) suggests that threshold exceedance captures local thermal accumulation more independently of site-level fixed effects. GEE-based correction substantially reduced the apparent predictive performance of DHW (AUC: 0.758 uncorrected → 0.624 corrected), suggesting that previously reported DHW performance may have been partially inflated by within-site spatial autocorrelation.

To examine the correspondence between threshold exceedance and the bimodal bleaching structure documented in Section 3.2, we cross-tabulated site-year median bleaching domain (low/intermediate/high) against days above 30°C categories (n = 93 site-years from 21 sites; some sites were not surveyed in all years, Section 2.3). When SST never exceeded 30°C, all site-years fell in the low-bleaching domain (39/39, 100%). At 1–20 days above 30°C, site-years were split between low (55.0%) and intermediate (42.5%) domains, with only 2.5% reaching the high domain. At 21 or more days above 30°C, 71.4% of site-years were in the high-bleaching domain (χ^2^ = 80.94, df = 4, p < 0.001; note that site-years within the same site are not independent, so the p-value is indicative rather than exact; Supplementary Table S7). This site-level correspondence provides empirical support for the interpretation that the bimodal distribution and the predictive advantage of threshold exceedance logic reflect the same underlying state transition dynamics.

**1.4 Sensitivity analysis across absolute thresholds from 27°C to 32°C confirmed that the 30°C threshold adopted a priori from Sakai et al. (2019) was near-optimal for both the >0% and ≥50% response definitions (Supplementary Table S1). AUC values formed a plateau between 29.5°C and 30.5°C, declining sharply at lower and higher thresholds**.

## 1.5 4. Discussion

This study reports three descriptive findings from five years (2020–2024) of standardised coral bleaching monitoring across 26 sites and 585 survey points in the Ryukyu Archipelago and adjacent waters. First, bleaching prevalence is bimodally distributed, with a stable intermediate zone and inter-annual variation driven by redistribution between low and high domains. Second, the two major bleaching years in the dataset — 2022 and 2024 — are qualitatively distinct events that cannot be collapsed into a single “mass bleaching” category. Third, the relative performance of thermal stress metrics depends on the ecological severity of the response being predicted: threshold exceedance logic discriminates moderate-to-severe bleaching more effectively than cumulative DHW. These findings carry implications for how bleaching data are modelled, how bleaching events are classified, and how the design space of thermal stress metrics is explored.

### 1.5.1 4.1 Bimodality and threshold behaviour

The most striking structural feature of these data is the bimodal distribution of bleaching prevalence, confirmed independently by Hartigan’s dip test (all years p < 0.001; Table 4) and beta mixture modelling (ΔBIC = 9.0–113.9; Table 5). The intermediate range (20–80%) accounts for a remarkably stable 21–27% of observations across all five years (Table 3), while inter-annual variation is driven almost entirely by redistribution between the low (<20%) and high (≥80%) domains. If bleaching progressed as a continuous gradient in response to thermal stress, one would expect the intermediate zone to expand in severe years; instead, it remains fixed. This pattern is consistent with a threshold-mediated binary switch rather than a dose-response continuum.

Multiple independent lines of physiological evidence support threshold behaviour in the bleaching response. Pfab et al. (2024) modelled the bioenergetic dynamics of the coral–dinoflagellate symbiosis under thermal stress and showed that once heat-induced photodamage exceeds a critical level, the energy balance of the holobiont collapses and symbiont expulsion becomes self-reinforcing — an escalating positive feedback that predicts minimal bleaching below the threshold and rapid progression to near-total bleaching above it. This model is situated within the broader mechanistic framework synthesised by Helgoe et al. (2024), who integrated multiple bleaching pathways into a triggers–cascades–endpoints architecture: diverse initial triggers (thermal, light, oxidative) converge on shared intracellular cascades that, once initiated, proceed largely independently of the triggering stimulus — a convergence structure that itself predicts threshold-like dynamics at the organismal level. Complementary to this ROS-centred cascade, Wooldridge (2009) proposed that thermal stress disrupts the CO -concentrating mechanism of Symbiodiniaceae, leading to photorespiratory production of glycolate and toxic accumulation of hydrogen peroxide — a pathway that likewise predicts a sharp transition once the concentrating mechanism fails. These models differ in the specific rate-limiting step but converge on the prediction that bleaching initiation is governed by a threshold rather than by a graded dose-response. Our distributional data are consistent with this convergent prediction operating at the site level.

However, the distributional pattern alone cannot adjudicate between physiological threshold mechanisms and alternative explanations for bimodality. Spatial autocorrelation of environmental conditions could produce clustered bleaching outcomes that mimic threshold dynamics without requiring a physiological switch. Unmeasured covariates that vary at the site level (e.g., water flow regime, depth profile, species composition, symbiont community structure) could similarly generate bimodal outcomes through environmental filtering.

That the bimodal pattern is driven by between-site divergence rather than within-site variability (Section 3.5; ICC = 0.76–0.89) is necessary for but not sufficient to establish a physiological threshold mechanism. The bimodality is a robust descriptive finding; its causal attribution remains open.

That entire sites fell cleanly into either the low or the high domain in 2024 (Section 3.5) indicates that whatever determines bleaching outcome operates at the site scale or above. This spatial polarisation implies that site-level environmental factors — not captured by satellite SST alone — filter the bleaching response, whether through modulating a physiological threshold or through independent environmental determination.

To our knowledge, this bimodal pattern has not been characterised in the coral bleaching literature as a persistent distributional feature across repeated annual surveys. Most large-scale analyses operate on bleaching occurrence as a binary response (Sully et al. 2019) or colony-level scores and cannot resolve the continuous site-level prevalence structure documented here. The bimodal structure implies that zero-one inflated beta regression (Ospina & Ferrari 2012) is the appropriate modelling framework for these data, a point we return to in the Limitations.

If bimodality reflects a genuine state transition, it carries a structural implication for thermal stress prediction. The cumulative logic of DHW — in which bleaching intensity scales with integrated heat exposure — is designed to predict a graded dose-response. But if the ecologically relevant outcome is a binary switch between unbleached and bleached states, then what determines reef fate is whether a threshold is crossed, not how much heat is accumulated beyond it. The metric comparison in Section 3.7, where threshold exceedance logic outperforms DHW for the state transition (≥50% bleaching), is consistent with this interpretation.

Growth-form-specific analyses indicate that bimodality is not an artefact of Acropora dominance at particular sites: Hartigan’s dip test rejected unimodality in all three growth-form categories (Acropora-dominant: D = 0.058; Mixed: D = 0.053; Other: D = 0.036; all p < 0.001; Fig. S1, Table S3). The persistence of bimodality across growth forms is more parsimoniously explained by site-level environmental filtering than by species composition effects alone, although unmeasured species-level differences in thermal tolerance may contribute at some sites. Nor is bimodality an artefact of observer rounding: the 135 unique values in the dataset, recorded at approximately 0.5% resolution, indicate that observers did not round to coarse categories, and bimodality was confirmed in the intermediate range (0 < bleaching < 100%) where boundary inflation at 0% and 100% is excluded.

### 1.5.2 4.2 Two anomalous years: 2022 versus 2024

Both 2022 and 2024 were major bleaching years, but their distributional signatures diverge in ways that carry interpretive consequences. In 2022, bleaching was positively skewed (skewness = 0.413): most sites experienced low-to-moderate bleaching, with a tail of severe cases. In 2024, the distribution was approximately symmetric (skewness = −0.100), the median reached 60.0%, and the high-bleaching component of the beta mixture model dominated (w = 0.616 vs 0.497). We characterise 2022 as a “partial mass bleaching” — selective, with a subset of sites severely affected — and 2024 as a “comprehensive mass bleaching” that engulfed the monitoring network.

This distinction matters because “mass bleaching” is routinely treated as a monolithic category. Existing classification schemes operate at different scales and capture different attributes: Spady et al. (2026) define global mass bleaching events by geographic extent across ocean basins, while Hughes et al. (2017) categorised aerial survey severity using an ordinal scale at the individual reef level. Neither framework captures the distributional structure of prevalence across a monitoring network — the within-event heterogeneity that distinguishes selective from comprehensive bleaching. We do not propose a typology from two events, but demonstrate that distributional parameters — specifically, mixture component weights and skewness — provide a descriptive vocabulary for this within-event variation that scalar summaries and ordinal categories alike obscure. Whether partial and comprehensive bleaching represent discrete categories or endpoints of a continuum requires comparison across regions and monitoring programs.

Despite its greater extent, the 2024 event did not produce proportionally greater mortality at the time of survey (mean 18.7% vs 15.2% in 2022). Mellin et al. (2019) documented a similar decoupling between bleaching severity and coral cover change at continental scales on the GBR, suggesting that more extensive bleaching does not necessarily translate to greater structural loss. Recurrent bleaching can also progressively restructure reef assemblages through differential mortality of heat-sensitive taxa (Hughes et al. 2018b), a process that may underlie the qualitative shift observed between the 2022 and 2024 events in this study.

As of our literature search (Web of Science, Scopus, Google Scholar; 6 February 2026), no peer-reviewed primary study has documented the 2024 bleaching event in Japan. Afzal et al. (2024) reported the 2022 event at Sekisei Lagoon. The present study provides the first systematic comparison of both events under identical monitoring protocols.

### 1.5.3 4.3 Bleaching, mortality, and the limits of annual monitoring

The relationship between bleaching prevalence and mortality is non-linear (Table 7). Mortality escalates sharply above 80% bleaching — mean 30.9%, with 96.4% of sites recording some mortality — while the 50–80% band shows mean mortality of only 13.2%, a 2.3-fold step across a single prevalence band. This threshold-like escalation in mortality is consistent with Brodnicke et al. (2019), who documented that severely bleached tabular *Acropora* colonies experienced markedly increased disease susceptibility and whole-colony mortality.

The most notable finding, however, is that even among sites with ≥80% bleaching, the median mortality was only 25.0% — that is, at least half of all severely bleached sites had experienced mortality of 25% or less at the time of survey. As noted in Section 2.2, surveys are conducted 1–3 months after the bleaching peak, and Sakai et al. (2019) documented post-bleaching mortality progressing over six months with four-fold variation within the same genus (*Acropora*). The mortality values in Table 7 therefore represent a lower bound, and the true extent of post-bleaching mortality in 2022 and 2024 remains unknown.

Spot-level tracking reveals that bleaching prevalence declined sharply from 2022 to 2023, but annual snapshots cannot distinguish genuine recovery from recruit replacement or delayed mortality (Morais et al. 2021). Back-to-back tracking across 2022 and 2024 shows that 32.0% of survey points experienced ≥50% bleaching in both years, while 9.4% transitioned from unaffected to severely bleached — whether this reflects cumulative damage or loss of a site-level protective factor cannot be resolved from prevalence data alone.

### 1.5.4 4.4 Metric structure, response definition, and the design space of thermal stress metrics

The predictive comparison reveals a consistent structural advantage of threshold exceedance logic across all bleaching severity levels. For detection-limit bleaching (>0%), days above 30°C (AUC 0.830) outperformed DHW (AUC 0.633). For moderate bleaching (≥50%), the advantage was greatest (AUC 0.877 vs 0.624; p < 0.001). For severe bleaching (≥80%), days above 30°C maintained a substantial advantage (AUC 0.895 vs 0.696). The threshold metric’s AUC jump from 0.830 to 0.877 is expected given the binary dynamics of the bimodal distribution: the ecologically significant state transition occurs above, not at, the detection limit.

This reframes the question from “which metric is better?” to “which metric is appropriate for which ecological question?” DHW integrates cumulative heat exposure relative to the local Maximum Monthly Mean, capturing a site’s departure from its thermal baseline. Threshold exceedance logic is better suited to predicting whether a site will undergo the state transition that determines reef trajectory.

However, a fundamental limitation of the days-above-30°C metric must be acknowledged. The monitoring network spans 9° of latitude, and the 30°C threshold has an inherent geographic bias: at high-latitude sites (e.g., Kushimoto, 33.5°N), summer SST rarely reaches 30°C even in anomalously warm years, yet bleaching does occur. DHW, by contrast, uses site-specific MMM-relative anomalies that automatically adjust for latitudinal differences in baseline temperature — a structural advantage for detecting thermal anomalies across latitudinal gradients, even though days above 30°C nonetheless achieved significantly higher AUC at all response thresholds. Sensitivity analysis excluding two high-latitude sites where SST never reached 30°C (sites 1 and 19; Supplementary Table S5) yielded AUC values of 0.873 (days above 30°C) and 0.630 (DHW) at the ≥50% threshold, confirming that the metric advantage is not driven by true-negative inflation at these sites. Similarly, replacing the MUR-derived DHW with NOAA CoralTemp DHW (Supplementary Table S6) did not change the ranking: CoralTemp DHW AUC (0.633 at ≥50%) was comparable to MUR DHW (0.624), confirming that the result reflects the structural difference between cumulative and threshold logic rather than idiosyncrasies of our DHW implementation.

The question raised in the Introduction — whether cumulative heat dose is the most appropriate predictor category — receives a qualified answer from these data. For ecologically significant bleaching (≥50% prevalence), threshold exceedance logic is the more appropriate predictor category; for detection-limit bleaching, the distinction is less clear-cut. The question is not binary but severity-dependent.

This analysis points toward a natural direction for metric development. The logical endpoint is a site-specific MMM-relative threshold exceedance count — a metric that combines DHW’s climatological relativism with threshold exceedance logic’s binary structure, asking: on how many days did SST exceed MMM + Δ°C, where Δ is optimised regionally? This formulation subsumes both DHW and absolute threshold counting as special cases. Cheung et al. (2025) demonstrated through interpretable machine learning on the GBR that non-linear interactions between heat stress and environmental covariates resolve bleaching outcomes that accumulated heat stress alone cannot predict, reinforcing the case for moving beyond single-variable cumulative metrics. Prior DHW optimisation studies have focused on internal parameters — accumulation window length (Lachs et al. 2021), cutoff temperature and regional calibration (Whitaker & DeCarlo 2024). Our findings raise a structural question: whether cumulative integration is the appropriate predictor category for ecologically significant bleaching, and whether the transition from cumulative to threshold logic entails a simultaneous transition from absolute to relative thresholds.

### 1.5.5 4.5 Limitations

Six limitations qualify these findings.

First, surveys are annual autumn snapshots (Section 2.2). The mortality values reported here are conservative estimates that do not capture the full delayed mortality process.

Second, vertical and lateral environmental data — water flow, turbidity, light environment, dissolved oxygen, temperature stratification — are absent from the monitoring protocol. The site-level environmental factors that produce the spatial polarisation documented in Section 4.1 remain unidentified. Without direct environmental measurements, the environmental heterogeneity hypothesis is observationally motivated but untested.

Third, the 2021 Yaeyama region presents a pattern consistent with non-random logger deployment: logger sites recorded near-zero bleaching while non-logger sites recorded 100% bleaching (Fisher’s exact test, p 0.001). The monitoring protocol directs loggers to locations where temperature fluctuations are pronounced, criteria that may favour microhabitats with historically persistent coral communities (Little & Rubin 2019; Bowler et al. 2025). If logger sites systematically sample sheltered microhabitats, satellite-logger discrepancies may overestimate satellite bias — a structural limitation that warrants systematic investigation (Supplementary Table S2).

Fourth, species-level resolution is lacking. Bleaching prevalence is a species-agnostic aggregate, and the mortality decoupling documented in Section 4.3 is likely driven by species, genotype, and symbiont differences that site-level surveys cannot resolve. Species composition provides an alternative explanation for bimodality: sites dominated by thermally sensitive genera (e.g., *Acropora*) may cluster in the high-bleaching domain independent of any physiological threshold mechanism. The Monitoring Site 1000 protocol records genus-level composition at a subset of sites; systematic analysis of composition effects on the distributional structure is deferred to subsequent work.

Fifth, spatial autocorrelation is not accounted for. Survey points within sites are spatially proximate and not independent; mixed-effects models with site as a random effect are required for inferential analyses.

Sixth, metric performance is sensitive to the binary response threshold, though GEE-based tests confirmed statistical significance at all three thresholds (Section 3.7). The choice of exchangeable working correlation structure assumes constant within-site correlation; alternative structures (e.g., unstructured, AR(1)) were not compared and may yield different standard errors for the AUC estimates. Any binary classification of a continuous variable entails information loss; zero-one inflated beta regression (Section 4.1) would allow modelling the full distributional response without dichotomisation.

The bimodal distributional structure implies that standard linear or beta regression models are misspecified for these data; zero-one inflated beta regression (Ospina & Ferrari 2012) is the statistically appropriate framework, and its implementation is a priority for subsequent work.

## 1.5.6 Conclusions

### Established

(1) Bleaching prevalence in the Ryukyu Archipelago is bimodally distributed, with a stable intermediate zone — a pattern consistent with threshold-mediated state transitions rather than continuous dose-response. (2) The 2022 and 2024 bleaching events are qualitatively distinct: partial versus comprehensive mass bleaching, distinguishable by distributional parameters that provide a descriptive vocabulary beyond scalar summaries. (3) The predictive performance of thermal stress metrics depends on response definition: threshold exceedance logic discriminates bleaching more effectively than cumulative DHW across all severity levels (GEE cluster bootstrap Wald test: >0%, p = 0.007; ≥50%, p < 0.001; ≥80%, p = 0.002) — indicating that metric structure and ecological severity threshold are inseparable design considerations.

### Open

(1) The environmental factors that produce site-level spatial polarisation. (2) The generality and magnitude of logger deployment bias across the monitoring network. (3) Whether the bimodality reflects physiological thresholds, environmental filtering, or their interaction. (4) The full extent of post-bleaching mortality beyond the autumn survey window. (5) Optimal site-specific MMM-relative threshold values across the latitudinal range of the monitoring network. (6) Whether a site-specific MMM-relative threshold exceedance count — combining DHW’s climatological relativism with threshold exceedance logic — improves predictive accuracy across response definitions and latitudinal gradients.

## 1.7 Acknowledgements

We thank the Biodiversity Center of Japan, Ministry of Environment, for providing access to the Monitoring Site 1000 coral reef survey data (approval number Kansei-Tahatsu No. 2602271). Data citation: Ministry of Environment, Japan. Monitoring Site 1000 Coral Reef Survey (FY2020-2024). This research received no specific grant from any funding agency in the public, commercial, or not-for-profit sectors.

## 1.8 Author contributions

HF conceived the study, conducted all analyses, and wrote the manuscript.

## 1.10 Data availability

Processed data and analysis code are available at https://github.com/hirokifukui/marine-obs. The raw Monitoring Site 1000 data are administered by the Biodiversity Center of Japan, Ministry of Environment, and available upon application (https://www.biodic.go.jp/moni1000/).

## 1.12 Data and Code Availability

All analysis code and processed data are available at https://github.com/hirokifukui/marine-obs. Raw monitoring data were provided by the Biodiversity Center of Japan, Ministry of the Environment (approval number Kansei-Tahatsu No. 2602271).

## 1.13 Supplementary Material (additional tables)

### 1.13.1 Table S1. AUC values for days above threshold temperatures (27–32°C) in predicting bleaching prevalence ≥ 50%

Sensitivity analysis of absolute temperature threshold selection. Days exceeding each threshold were calculated from MUR SST daily data across all survey-point observations (n = 2,288; 26 sites). AUC values show a plateau between 29.5°C and 30.5°C, supporting the a priori selection of 30°C used in the main analysis.

| Threshold (°C) | AUC |
| --- | --- |
| 27.0 | 0.608 |
| 27.5 | 0.599 |
| 28.0 | 0.599 |
| 28.5 | 0.627 |
| 29.0 | 0.680 |
| 29.5 | 0.754 |
| 30.0 | 0.818 |
| 30.5 | 0.821 |
| 31.0 | 0.757 |
| 31.5 | 0.652 |
| 32.0 | 0.577 |

Response variable: bleaching prevalence ≥ 50% (binary). MUR SST (NASA, 1 km resolution).

### 1.13.2 Table S2. Sensitivity analysis: exclusion of Yaeyama 2021 observations

As a sensitivity analysis, we excluded observations from Yaeyama sites (site11, site12) in 2021, where temperature logger placement may have been influenced by prior bleaching history (n = 4 observations excluded; all with bleaching prevalence = 0%). AUC values were essentially unchanged (Table S2), confirming that the deployment bias in this subset does not materially affect the main conclusions.

| Metric | Full dataset (N = 186) | Excl. Yaeyama 2021 (N = 182) | $\Delta$ AUC |
| --- | --- | --- | --- |
| Days above 30°C | 0.875 | 0.879 | +0.004 |
| DHW | 0.638 | 0.646 | +0.008 |

### 1.13.3 Table S3. Hartigan’s dip test for bimodality by coral growth-form category

Bleaching prevalence was pooled across all five fiscal years and stratified by dominant growth form at each survey point. Dip test applied to the full range (0–100%). All three categories rejected unimodality, indicating that bimodality is not an artefact of taxonomic composition at specific sites.

| Growth-form category | n | Dip statistic (D) | p |
| --- | --- | --- | --- |
| Acropora-dominant | 917 | 0.058 | <0.001 |
| Mixed | 814 | 0.053 | <0.001 |
| Other | 557 | 0.036 | <0.001 |

### 1.13.4 Table S4. Bootstrap 95% confidence intervals for GEE-based AUC values

Cluster bootstrap (10,000 iterations, site-level resampling, seed = 42) 95% confidence intervals for each metric’s AUC at three bleaching response thresholds. GEE with exchangeable correlation structure, 21 sites.

| Response threshold | Metric | AUC | 95% CI |
| --- | --- | --- | --- |
| >0% (any bleaching) | Days above 30°C | 0.830 | [0.722, 0.918] |
|  | DHW | 0.633 | [0.552, 0.734] |
| $\geq 50\%$ (moderate–severe) | Days above 30°C | 0.877 | [0.760, 0.964] |
|  | DHW | 0.624 | [0.537, 0.740] |
| $\geq 80\%$ (severe) | Days above 30°C | 0.895 | [0.775, 0.965] |
|  | DHW | 0.696 | [0.616, 0.793] |

### 1.13.5 Table S5. Sensitivity analysis: exclusion of high-latitude sites where SST never reached 30°C

Two sites (site 1, 30.46°N; site 19, 34.98°N) never recorded any days above 30°C across all five fiscal years. AUC values after excluding these sites (n = 175 site-year observations, 19 sites) confirm that the predictive advantage of threshold exceedance is not driven by true-negative inflation.

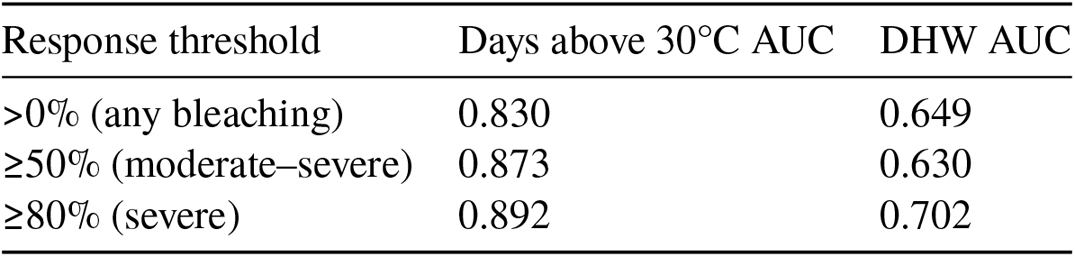

| Response threshold | Days above 30°C AUC | DHW AUC |
| --- | --- | --- |
| >0% (any bleaching) | 0.830 | 0.649 |
| ≥50% (moderate–severe) | 0.873 | 0.630 |
| ≥80% (severe) | 0.892 | 0.702 |

### 1.13.6 Table S6. Comparison of MUR-derived DHW and NOAA CoralTemp DHW

To address the concern that results might reflect idiosyncrasies of the MUR-derived DHW implementation, we repeated the AUC comparison using DHW computed from the NOAA CoralTemp SST product (same algorithm: MMM+0°C cutoff, 84-day window). CoralTemp DHW AUC values were comparable to MUR DHW, confirming that the structural advantage of threshold exceedance logic is robust to the choice of SST product.

| Response threshold | Days above 30°C AUC | CoralTemp DHW AUC | MUR DHW AUC |
| --- | --- | --- | --- |
| >0% (any bleaching) | 0.830 | 0.635 | 0.633 |
| ≥50% (moderate–severe) | 0.877 | 0.633 | 0.624 |
| ≥80% (severe) | 0.895 | 0.728 | 0.696 |

### 1.13.7 Table S7. Cross-tabulation of days above 30°C categories and bleaching domain

Site-year median bleaching prevalence was classified into three domains (Low: <20%, Intermediate: 20–80%, High: ≥80%). Days above 30°C were categorised as 0 days, 1–20 days, or 21+ days. n = 93 site-years (21 sites × up to 5 years). χ^2^ = 80.94, df = 4, p < 0.001.

#### Frequency

Frequency:
| Days above 30°C | Low (<20%) | Intermediate (20–80%) | High (≥80%) | Total |
| --- | --- | --- | --- | --- |
| 0 days | 39 | 0 | 0 | 39 |
| 1–20 days | 22 | 17 | 1 | 40 |
| 21+ days | 1 | 3 | 10 | 14 |
| Total | 62 | 20 | 11 | 93 |

#### Row percentages

Row percentages:
| Days above 30°C | Low (<20%) | Intermediate (20–80%) | High (≥80%) |
| --- | --- | --- | --- |
| 0 days | 100.0% | 0.0% | 0.0% |
| 1–20 days | 55.0% | 42.5% | 2.5% |
| 21+ days | 7.1% | 21.4% | 71.4% |

## References

Afzal MS, Udo T, Ueno M, Nakamura T (2024) Mass coral bleaching and mortality associated with high sea surface temperatures in the summer of 2022 in Sekisei Lagoon, Okinawa, Japan. Galaxea, Journal of Coral Reef Studies 26:20–26. doi:10.3755/galaxea.g26n-5

Bowler DE, Boyd RJ, Callaghan CT, Robinson RA, Isaac NJB, Pocock MJO (2025) Treating gaps and biases in biodiversity data as a missing data problem. Biological Reviews 100(1):50–67. doi:10.1111/brv.13127

Brodnicke OB, Bourne DG, Heron SF, Pears RJ, Stella JS, Smith HA, Willis BL (2019) Unravelling the links between heat stress, bleaching and disease: fate of tabular corals following a combined disease and bleaching event. Coral Reefs 38:591–603. doi:10.1007/s00338-019-01813-9

Cheung MWM, Chaloupka M, Hock K, Mumby PJ (2025) Moving beyond temperature metrics in coral bleaching prediction using interpretable machine learning. Global Ecology and Biogeography 34:e70105. doi:10.1111/geb.70105

DeLong ER, DeLong DM, Clarke-Pearson DL (1988) Comparing the areas under two or more correlated receiver operating characteristic curves: a nonparametric approach. Biometrics 44(3):837–845.

Hartigan JA, Hartigan PM (1985) The dip test of unimodality. Annals of Statistics 13(1):70–84.

Helgoe J, Davy SK, Weis VM, Rodriguez-Lanetty M (2024) Triggers, cascades, and endpoints: connecting the dots of coral bleaching mechanisms. Biological Reviews 99(3):715–752. doi:10.1111/brv.13042

Hughes TP, Kerry JT, Álvarez-Noriega M, Álvarez-Romero JG, Anderson KD, Baird AH, Babcock RC, Beger M, Bellwood DR, Berkelmans R, Bridge TC, Butler IR, Byrne M, Cantin NE, Comeau S, Connolly SR, Cumming GS, Dalton SJ, Diaz-Pulido G, Eakin CM, Figueira WF, Gilmour JP, Harrison HB, Heron SF, Hoey AS, Hobbs J-PA, Hoogenboom MO, Kennedy EV, Kuo C-Y, Lough JM, Lowe RJ, Liu G, McCulloch MT, Malcolm HA, McWilliam MJ, Pandolfi JM, Pears RJ, Pratchett MS, Schoepf V, Simpson T, Skirving WJ, Sommer B, Torda G, Wachenfeld DR, Willis BL, Wilson SK (2017) Global warming and recurrent mass bleaching of corals. Nature 543:373–377. doi:10.1038/nature21707

Hughes TP, Anderson KD, Connolly SR, Heron SF, Kerry JT, Lough JM, Baird AH, Baum JK, Berumen ML, Bridge TC, Claar DC, Eakin CM, Gilmour JP, Graham NAJ, Harrison HB, Hobbs J-PA, Hoey AS, Hoogenboom M, Lowe RJ, McCulloch MT, Pandolfi JM, Pratchett MS, Schoepf V, Torda G, Wilson SK (2018a) Spatial and temporal patterns of mass bleaching of corals in the Anthropocene. Science 359:80–83. doi:10.1126/science.aan8048

Hughes TP, Kerry JT, Baird AH, Connolly SR, Dietzel A, Eakin CM, Heron SF, Hoey AS, Hoogenboom MO, Liu G, McWilliam MJ, Pears RJ, Pratchett MS, Skirving WJ, Stella JS, Torda G (2018b) Global warming transforms coral reef assemblages. Nature 556:492–496. doi:10.1038/s41586-018-0041-2

JPL MUR MEaSUREs Project (2015) GHRSST Level 4 MUR Global Foundation Sea Surface Temperature Analysis (v4.1). doi:10.5067/GHGMR-4FJ04

Kass RE, Raftery AE (1995) Bayes factors. Journal of the American Statistical Association 90(430):773–795. doi:10.1080/01621459.1995.10476572

Kawagoe S (2017) Mass coral bleaching in 2016 reported by the Monitoring sites 1000 project. Journal of Japanese Coral Reef Society 19:21–28. [in Japanese]

Kimura T, Chou LM, Huang D, Tun K, Goh E (eds) (2022) Status and Trends of East Asian Coral Reefs: 1983–2019. Global Coral Reef Monitoring Network (GCRMN) / International Coral Reef Initiative (ICRI).

Lachs L, Bythell JC, East HK, Edwards AJ, Mumby PJ, Skirving WJ, Spady BL, Guest JR (2021) Fine-tuning heat stress algorithms to optimise global predictions of mass coral bleaching. Remote Sensing 13(14):2677. doi:10.3390/rs13142677

Little RJA, Rubin DB (2019) Statistical Analysis with Missing Data, 3rd ed. John Wiley & Sons.

Liu G, Heron SF, Eakin CM, Muller-Karger FE, Vega-Rodriguez M, Guild LS, De La Cour JL, Geiger EF, Skirving WJ, Burgess TFR, Strong AE, Harris A, Maturi E, Ignatov A, Sapper J, Li J, Lynds S (2014) Reef-scale thermal stress monitoring of coral ecosystems: new 5-km global products from NOAA Coral Reef Watch. Remote Sensing 6(11):11579–11606. doi:10.3390/rs61111579

Mellin C, Matthews S, Anthony KRN, Brown SC, Caley MJ, Johns KA, Osborne K, Puotinen M, Thompson A, Wolff NH, Fordham DA, MacNeil MA (2019) Spatial resilience of the Great Barrier Reef under cumulative disturbance impacts. Global Change Biology 25:2431–2445. doi:10.1111/gcb.14625

Ministry of Environment, Japan (2024) Monitoring Site 1000 Coral Reef Survey Manual. https://www.biodic.go.jp/moni1000/mual/coral_manual.pdf

Morais J, Morais RA, Tebbett SB, Pratchett MS, Bellwood DR (2021) Dangerous demographics in post-bleach corals reveal boom-bust versus protracted declines. Scientific Reports 11:18787. doi:10.1038/s41598-021-98239-7

Ospina R, Ferrari SLP (2012) A general class of zero-or-one inflated beta regression models. Computational Statistics & Data Analysis 56(6):1609–1623. doi:10.1016/j.csda.2011.10.005

Pfab F, Detmer AR, Moeller HV, Nisbet RM, Putnam HM, Cunning R (2024) Heat stress and bleaching in corals: a bioenergetic model. Coral Reefs 43:1627–1645. doi:10.1007/s00338-024-02561-1

Sakai K, Singh T, Iguchi A (2019) Bleaching and post-bleaching mortality of Acropora corals on a heat-susceptible reef in 2016. PeerJ 7:e8138. doi:10.7717/peerj.8138

Scott DW (1992) Multivariate Density Estimation: Theory, Practice, and Visualization. John Wiley & Sons.

Spady BL, Skirving WJ, De La Cour JL, Geiger EF, Liu G, Hoegh-Guldberg O, Norrie A, Heron SF, Pomeroy MW, Kolodziej G, Brown KT, Manzello DP (2026) The 4th global coral bleaching event: ushering in an era of near-annual bleaching. Coral Reefs (advance online publication). doi:10.1007/s00338-025-02810-x

Sully S, Burkepile DE, Donovan MK, Hodgson G, van Woesik R (2019) A global analysis of coral bleaching over the past two decades. Nature Communications 10:1264. doi:10.1038/s41467-019-09238-2

Whitaker H, DeCarlo TM (2024) Re(de)fining degree-heating week: coral bleaching variability necessitates regional and temporal optimization of global forecast model stress metrics. Coral Reefs 43:969–984. doi:10.1007/s00338-024-02512-w

Wooldridge SA (2009) A new conceptual model for the warm-water breakdown of the coral–algae endosymbiosis. Marine and Freshwater Research 60:483–496. doi:10.1071/MF08251

