## Supplementary figures and images for "Bimodal distribution of coral bleaching prevalence is consistent with state transition dynamics in thermal stress response: five years of standardised monitoring in Japan"

### Supplementary Figure S1

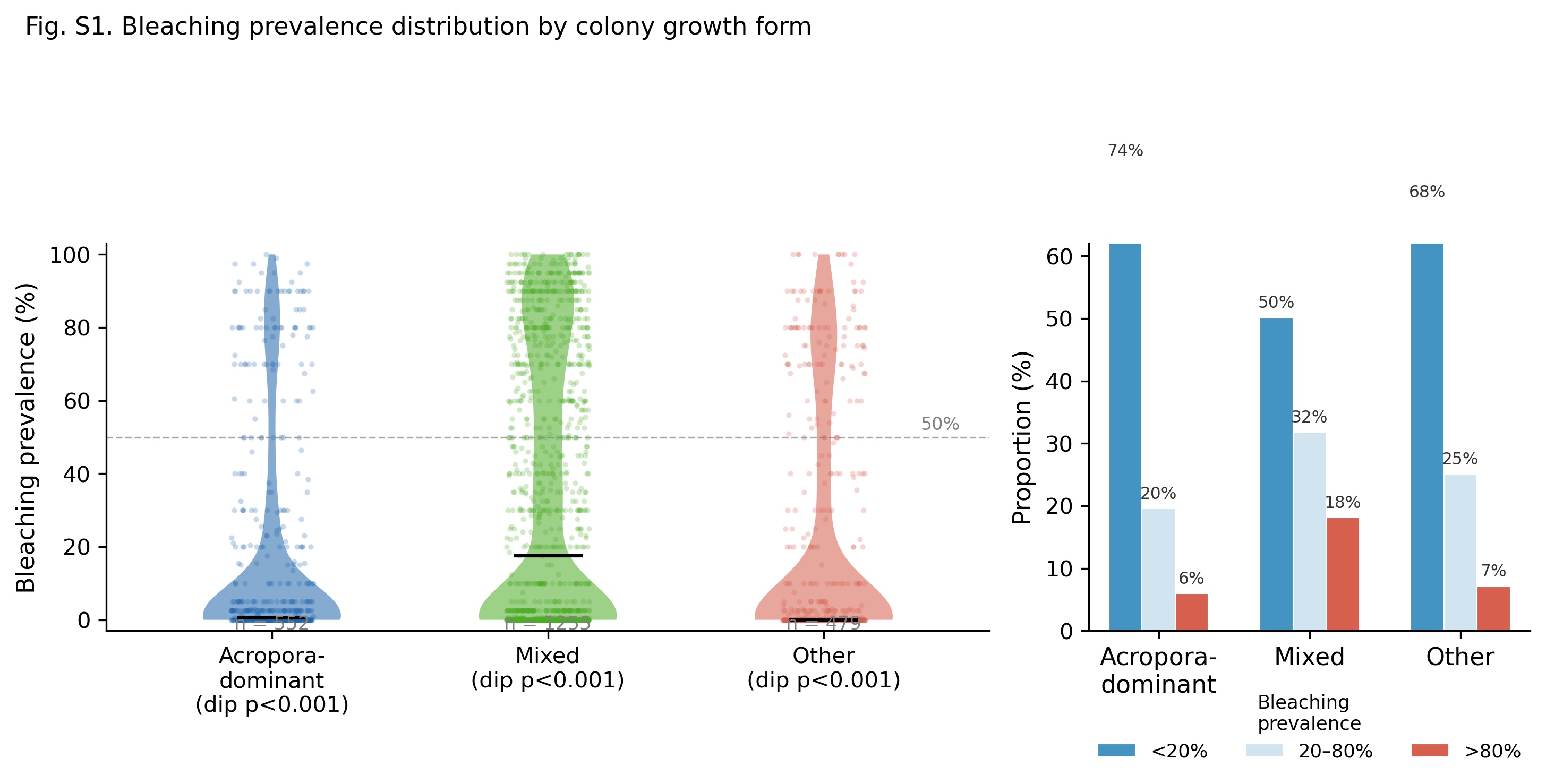
